# A causal role of the right superior temporal sulcus in emotion recognition from biological motion

**DOI:** 10.1101/079756

**Authors:** Rochelle A. Basil, Margaret L. Westwater, Martin Wiener, James C. Thompson

## Abstract

Understanding the emotions of others through nonverbal cues is critical for successful social interactions. The right posterior superior temporal sulcus (rpSTS) is one brain region thought to be key in the recognition of the mental states of others based on body language and facial expression. In the present study, we temporarily disrupted functional activity of the rpSTS by using continuous, theta-burst transcranial magnetic stimulation (cTBS) to test the hypothesis that the rpSTS plays a causal role in emotion recognition from body movements. Participants (N=23) received cTBS of the rpSTS, which was individually localized using fMRI, and a vertex control site. Before and after cTBS, we tested the ability of participants to identify emotions from point-light biological motion stimuli and a non-biological global motion identification task. Results revealed that the ability of participants to accurately identify emotional states from biological motion was reduced following cTBS stimulation to the rpSTS, but was unimpaired following vertex stimulation. Accuracy on the global motion tasks was unaffected by cTBS to either site. These results support the causal role of the rpSTS in decoding information about other’s emotional state from their body movements and gestures.

## Introduction

The dynamics and kinematics of body movements play a vital role in the expression and perception of emotion and other social cues. Indeed, the ability to detect, identify, and respond appropriately to these dynamic nonverbal cues is central to social competence (Mehrabian & Ferris, 1967; Rosenthal, Hall, DiMatteo, Rogers, & Archer, 1979). Neuroimaging evidence has long implicated the right posterior superior temporal sulcus (rpSTS) as a cortical region central to the perception of social cues from body movements (Bonda, Petrides, Ostry, & Evans, 1996; Grossman & Blake, 2002; Puce, Allison, Bentin, Gore, & McCarthy, 1998). This region shows a preference for dynamic bodies and faces over static (Pitcher, Dilks, Saxe, Triantafyllou, & Kanwisher, 2011; Puce & Perrett, 2003), and it is sensitive to the configuration of a moving body (Thompson, Clarke, Stewart, & Puce, 2005). While neuroimaging studies indicate that both right and left posterior STS can response during the perception of body movement (e.g., Saygin et al., 2007), there is evidence of a right hemisphere bias (Beauchamp, Lee, Haxby, & Martin, 2003; Grossman & Blake, 2002; M. Pavlova, Lutzenberger, Sokolov, & Birbaumer, 2004; David Pitcher, Duchaine, & Walsh, 2014). Greater fMRI response selectivity to point-light displays of biological motion in the rpSTS is also associated with larger, more complex social networks, implying that the coding of the movements of others in this region is important for social abilities (Dziura & Thompson, 2014). However, techniques such as cross-sectional fMRI are correlational and limited in their ability in establishing the causal role of a brain region in a perceptual or cognitive function. Here, we used fMRI-guided repetitive transcranial magnetic stimulation (rTMS) to examine if the rpSTS is causally involved in the coding of social cues conveyed by body movements.

Understanding the causal role of the rpSTS in the perception of dynamic social stimuli is important, as dysfunction of the processing of face and body in this region has been linked to social deficits in autism spectrum disorder (ASD) (Koldewyn, Whitney, & Rivera, 2011; M. A. Pavlova, 2012) and schizophrenia (J Kim, Norton, McBain, Ongur, & Chen, 2013; Jejoong Kim, Doop, Blake, & Park, 2005). More recently, decreased fMRI activity to point-light biological motion in rpSTS has been proposed as a neurobiomarker for ASD (Björnsdotter et al., 2016). Studies of brain structure also point to altered cortical thickness of the pSTS in neurodevelopmental disorders (Zilbovicius et al., 2006), and protracted maturation of this region has been associated with altered functional network differentiation in children and adolescents with ASD (Shih et al., 2011). Shih et al. (2011) postulate that atypical development of the pSTS could indicate impaired functional specificity of this region in individuals with ASD, and this might contribute to the inability of some with neurodevelopmental disorders to discriminate dynamic social cues, such as subtle shifts in body language, that convey the intentions or emotions of another. As such, increased understanding the role of the rpSTS in the perception and discrimination of social information from biological motion is important for future work with these clinical populations.

One previous study using rTMS indicated that disruption of the pSTS impairs the discrimination of point-light biological motion from background noise (Grossman, Battelli, & Pascual-Leone, 2005). A second study reported a trend for lower detection of biological motion from noise following rTMS to the left pSTS, but did not examine right pSTS (van Kemenade, Muggleton, Walsh, & Saygin, 2012). This work indicates that the STS is important for the integration of moving point-lights into a moving human figure from background noise motion. However, it remains possible that the role of the rpSTS does not extend beyond this function. To determine if pSTS function extends further to the coding and identification of actions, we examined the effects of disruption by continuous theta-burst rTMS (cTBS) on a point-light emotion discrimination task (Atkinson, Dittrich, Gemmell, & Young, 2004). We chose this task because the discrimination of emotions from gestures and body movements is a key social ability. These stimuli were presented without irrelevant noise motion. To account for the possibility that any cTBS effects were due to non-specific global motion discrimination, we compared performance on the emotion discrimination task to a non-biological global motion discrimination task. Furthermore, to account for individual differences in the precise location of the rpSTS region involved in processing biological motion, we used fMRI-guided neuronavigation of the TMS coil. It was hypothesized that cTBS targeted over the rpSTS would significantly impair both accuracy and reaction time to the emotion recognition task, relative to stimulation of a vertex control site. We did not anticipate cTBS stimulation to the rpSTS or vertex to impair performance on the global motion control task.

## Materials & Methods

### Participants

Twenty-three healthy, right-handed adults (13 female; *M*_*age*_ = 25.2 y; *SD* = 3.8) with normal or corrected-to-normal vision participated in the study. Exclusion criteria for both the fMRI and TMS sessions were: history of traumatic brain injury, diagnosis of a neurological disorder, ferromagnetic implants, or current use of psychotropic medication. Prior to participation, participants provided written informed consent and completed both MRI and TMS safety screening procedures. The study procedure and all relevant materials were approved by the George Mason University Human Subjects Review Board (HSRB). Participants were informed at each session that they were free to withdraw from the study at any time for any reason, and they received monetary compensation for their time.

### fMRI localizer: Identification of rpSTS target site

#### fMRI data acquisition and preprocessing

To ensure accurate targeting of rpSTS for the cTBS, we first collected blood oxygen level-dependent (BOLD) fMRI data as participants viewed a block design paradigm, consisting of five blocks of scrambled and intact 12-dot PLDs (Dziura & Thompson, 2014). All stimuli were obtained from the Carnegie Mellon University Graphics Lab Motion Capture Database (http://mocap.cs.cmu.edu). The 2-s videos were presented during data acquisition in blocks of 16 s per condition, alternating with a 12-s fixation block, using Neurobehavioral Systems Presentation software (Version 16.3; http://www.neurobs.com/). To ensure active viewing of the stimuli, participants were instructed to press a button with their right index finger upon seeing a stimulus repeat (1-back). Visual stimuli were presented on a rear projection screen that was viewed using a head coil-mounted mirror. Functional images were acquired on a 3T Siemens Allegra Magnetom using a quadrature headcoil. Each functional run comprised 172 gradient echo echo-planar images (GE EPI) (33 interleaved axial slices; TR/TE = 2000/30ms, flip angle = 70°, 64×64mm matrix, field of view = 192mm^2^). At the completion of the task, a T1-weighted, whole-head structural scan was acquired using a three-dimensional, magnetization-prepared, rapid-acquisition gradient echo (MPRAGE) pulse sequence (TR = 2300ms, TE = 3.37 ms, flip angle = 7, FOV = 1260 mm^2^, 256×256mm matrix, 1 mm slices with sagittal acquisition).

Functional data were analyzed using the fMRI Expert Analysis Tool (FEAT) within the FSL software package (version 6.0; fsl.fmrib.ox.ac.uk). Preprocessing involved correction for signal inhomogeneity, brain extraction, motion correction, linear registration to the T1 anatomical images, and spatial smoothing at 6mm FWHM. Intact motion and scrambled motion blocks, as well as head motion regressors, were entered into a linear regression at each voxel, using generalized least squares with a voxel-wise, temporally and spatially regularized autocorrelation model. A Gaussian-weighted running line smoother (100s FWHM) was also included in the model to account for drift. Individual subject regions of interest (ROIs) were created from peak voxels in the rpSTS exhibiting activation in the intact vs scrambled PLDs at p < 0.05 (uncorrected). For subject-specific localization of the rpSTS, ROIs were created in each subject's native space. The average MNI coordinates for the rpSTS was 56.57 (*4.90*), −43.48 (*5.76*), 12.87 (*9.81*). Two sample ROIs are shown in Figure 1.

**Figure 1:**
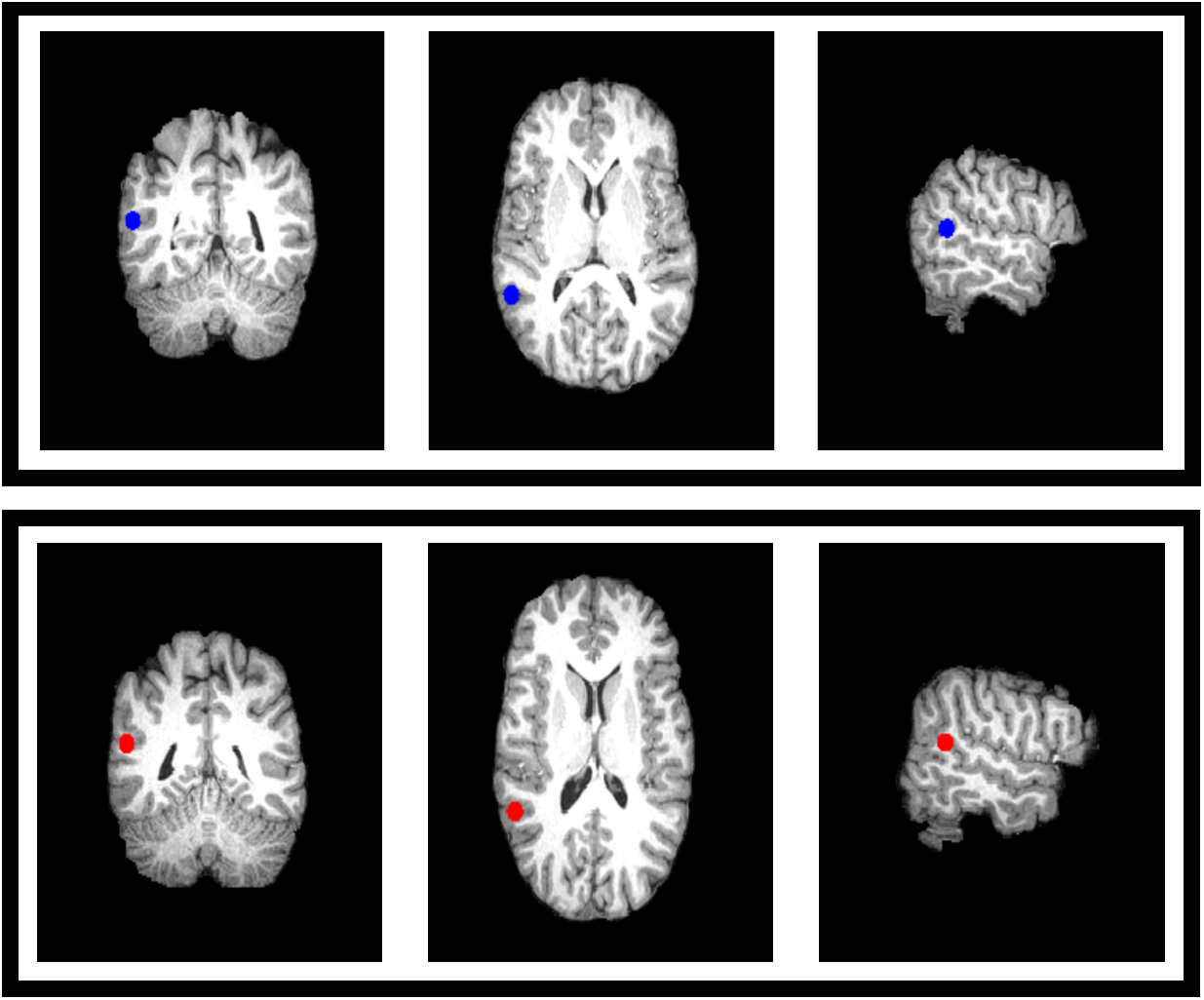
Example of two subjects rpSTS ROIs

### Theta Burst Stimulation of the rpSTS & Vertex

#### Apparatus and stimuli

Continuous theta burst stimulation of a 3 pulse burst at 50Hz, with a 5Hz inter-burst-interval was delivered in an uninterrupted train of 600 pulses, over each participant’s functionally-localized rpSTS and vertex, (as used by Pitcher et al. (2008), the midway point between the nasion and the inion, equidistant from the left and right intertragal notches). cTBS was delivered at 40% of maximal stimulator output (Huang, et al. 2005 by a Magstim Super Rapid+ Magnetic Stimulator with a 70mm diameter figure-of-eight, air-cooled coil (Magstim, UK), in conjunction with Brainsight Frameless Stereotaxic Software (Rogue Research, Montreal, Canada). This intensity was derived from previous studies (e.g., Borckardt et al., 2008). Participants were provided hearing protection for both stimulation sessions.

## Procedure

In a randomized and counterbalanced design, participants were presented with dynamic stimuli of PLDs, as well as global non-biological motion stimuli (dots moving at 20% coherence) in different directions (planar, radial, spiral) in separate alternating blocks. Three emotional states (angry, happy, and fearful) were selected, all of which are high on arousal but differ in terms of valence. Participants were instructed to identify the emotion (in the case of biological motion) or the type of motion (in the non-biological motion trials) using a keyboard as quickly and accurately as possible. Before the experiment began, participants completed a training session to familiarize themselves with both the stimuli and the button coding. In total, subjects completed four blocks of biological motion (two pre-TMS, two post-TMS) and four blocks of non-biological motion (two pre-TMS, two post-TMS) in a counterbalanced fashion. Each stimulus was presented for 1200 ms, and each block contained 48 trials (16 trials per condition). Each block was 3 minutes in duration. Stimuli were presented at a visual angle of 9.4° and a refresh rate of 60Hz. Each session lasted approximately 40 minutes. Stimuli were presented on a Dell E1911 19” Monitor with 1440x900 resolution and using Presentation Software (v. 17.2). Stimuli included in this experiment underwent pilot testing to ensure for a consistent level of baseline accuracy across both conditions. Stimuli were piloted with 12 participants who were not included in the main study.

## Results

The aim was to impair participants’ ability to identify emotional states from biological motion stimuli following cTBS over the rpSTS. As an active control, the vertex was stimulated. We did not anticipate a decrease in subjects’ ability to label emotional states after cTBS to the vertex. The main finding indicated that cTBS stimulation of the rpSTS reduced participants’ ability to accurately identify emotions, but such stimulation did not their ability to identify global motion types. There were no observed effects of either cTBS or stimulus type on reaction time.

Statistical analyses of task session data were completed in IBM SPSS Statistics (v.19). We conducted two, 2 × 2 repeated measures ANOVAs to examine the effect of cTBS site (rpSTS and vertex) and stimulus type (biological or non-biological) on accuracy and reaction time (RT).

### cTBS Stimulation on Accuracy

The 2 × 2 repeated measures ANOVA of accuracy showed a significant main effect of cTBS site, F(1,22) = 10.87, p = 0.003. A main effect of stimulus type also reached statistical significance F(1,22) = 8.50, p = 0.008.

Analyses also indicated a significant two-way interaction of TBS site and stimulus type, F(1,22) = 4.93, p = 0.037. These effects are shown in Figure 2. Post hoc paired samples t-tests revealed that the change in accuracy whencTBS was targeted over the rpSTS was significant between the two stimulus types (t(1,22) = −4.60, p < 0.001). Post hoc comparisons further demonstrated that participants’ ability to accurately identify emotions from biological motion was significantly impaired following cTBS to the rpSTS but not to the vertex (t(1,22) = −5.77, p < 0.001).

**Figure 2:**
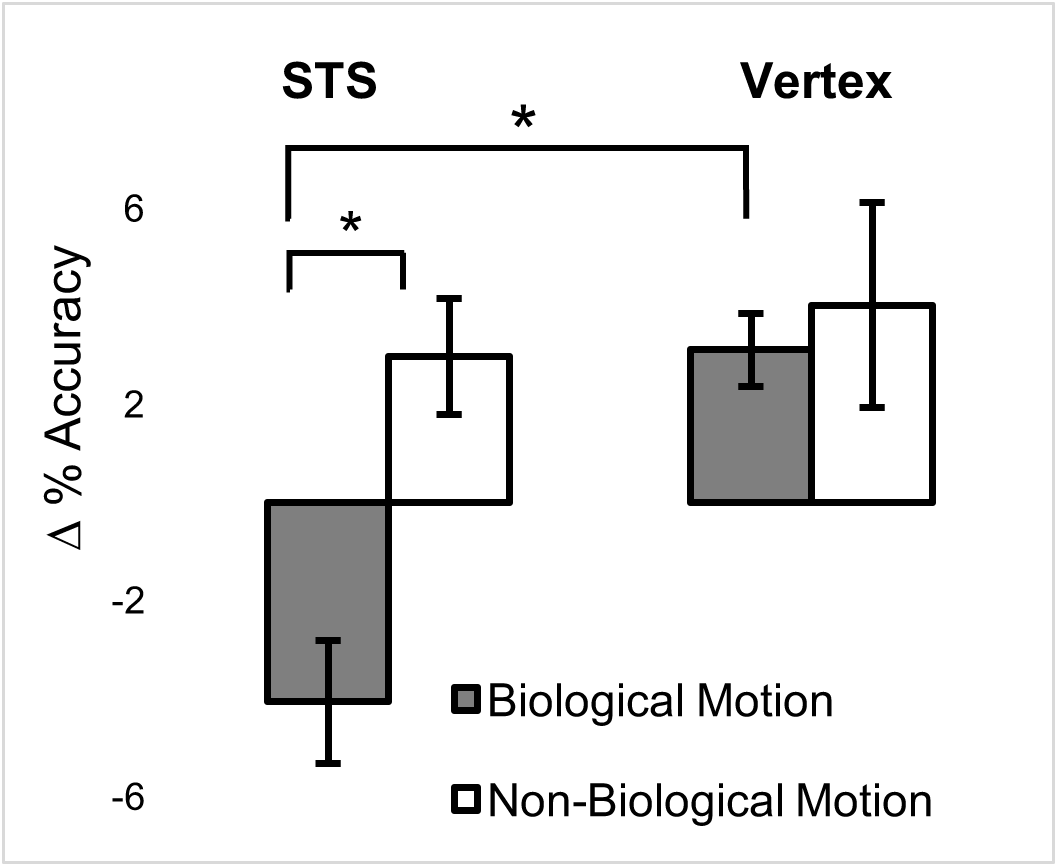
Mean differences in percentage change by condition (biological motion and non-biological motion) by TMS site (rpSTS and vertex). cTBS over rpSTS impaired emotional recognition through biological motion, but not non-biological motion. An asterisk denotes a significant (*p* < 0.001) difference. Error bars indicate SEs.

### cTBS Stimulation on Reaction Time

A 2 × 2 repeated measures ANOVA of RT did not demonstrate a main effect of cTBS site, F(1,22) = 0.37; p = 0.547, or stimulus type, F(1,22) = 2.89, p = 0.103, on RT. The two-way interaction of cTBS site and stimulus type was nonsignificant, F(1,22) = 0.00, p = 0.984. These findings are illustrated in Figure 3.

**Figure 3:**
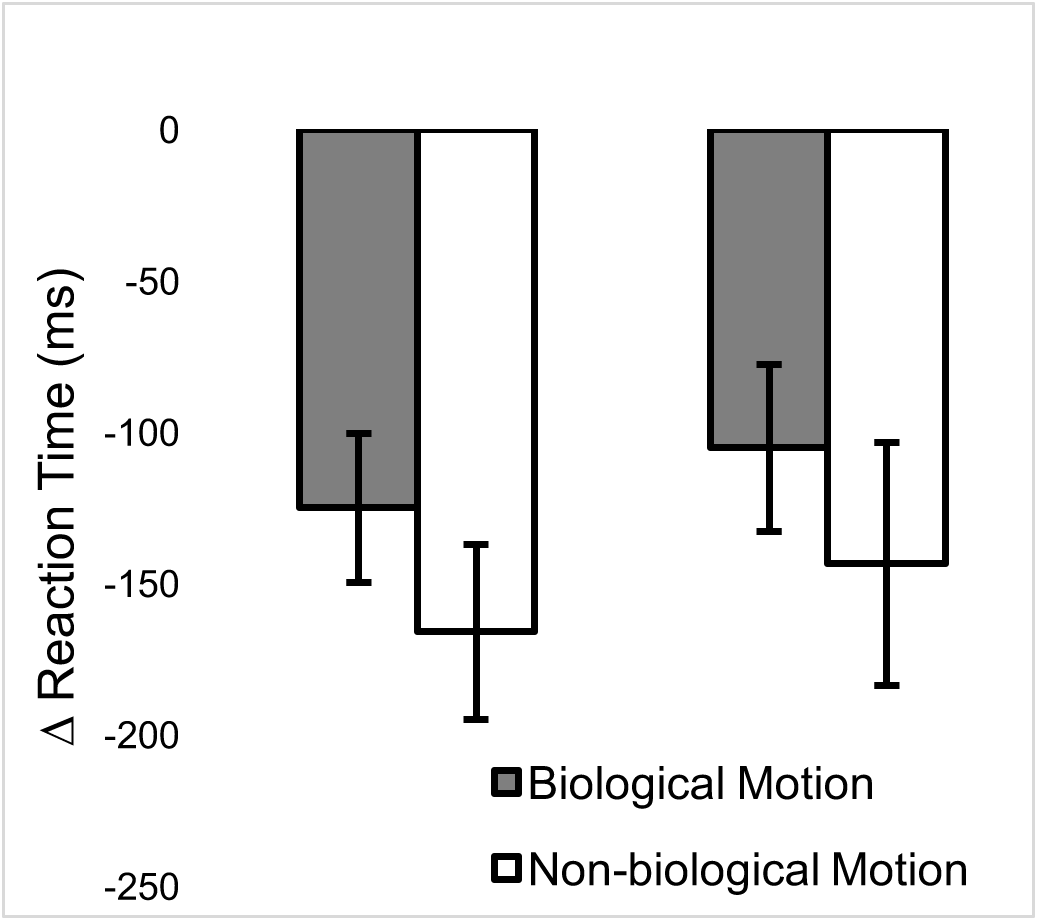
Mean differences in change in reaction time by condition (biological motion and non-biological motion) by TMS site (rpSTS and vertex). cTBS over rpSTS and vertex did not significantly decrease reaction time for either stimulus type. Error bars indicate SEs.

## Discussion

The present study found that disruption of the rpSTS using cTBS lead to selective impairment in the recognition of emotions conveyed by human movements. We used fMRI to target a rpSTS region in each participant, as this region has been previously implicated in thevisual processing of biological motion and in ‘social networks’ of the brain. We found that right STS-targeted cTBS reduced the accuracy of the identification of different emotional point-light stimuli. The detrimental effects of cTBS to this region did not extend to the recognition of non-biological motion, as we found no significant impairment in the identification of different global motion stimuli. The effect of cTBS to the rpSTS was limited to recognition accuracy, as reaction time was left unaffected. The findings of this study indicate that the rpSTS subserves the coding of dynamic social information, such as emotion, conveyed by the body movements of another person.

Previous work has shown that 1Hz rTMS to the pSTS region can impair discrimination of biological motion from noise (Grossman et al., 2005). Additional work has indicated that rTMS to the pSTS can decrease walking direction discrimination from biological motion while leaving facing orientation unaffected (Vangeneugden, Peelen, Tadin, & Battelli, 2014). Taken together, these studies underscore the importance of a sub-region of the pSTS to the detailed visual processing of biological motion; however, it remains unclear whether the causal role of this pSTS region extends to the coding of social information, such as emotion. It is possible that the pSTS serves as a high-level visual processor that is not central to the processing of social meaning from such stimuli. The selectivity of the fMRI response of the pSTS region to biological motion region does appear to be linked to social abilities (Pelphrey, Morris, & McCarthy, 2004; Saxe, Xiao, Kovacs, Perrett, & Kanwisher, 2004), and even the size and complexity of social networks (Dziura & Thompson, 2014), suggesting a more diffuse role in social cognition. More recently, an fMRI study showed that the pSTS region is involved in the processing of emotion conveyed by body movements (Goldberg, Christensen, Flash, Giese, & Malach, 2015). Together with the findings of the present study, which demonstrate a causal role in emotion recognition from biological motion, it seems that the visual coding of human movements represents one facet of a broader social cognitive role played by the pSTS region (Allison, Puce, & McCarthy, 2000).

Consistent with a broader social cognitive role of the rpSTS region, this region has also been considered by many as part of a face processing network (Haxby, Hoffman, & Gobbini, 2000, 2002; Hoffman & Haxby, 2000). The original model by Haxby and colleagues (2000) proposed that the rpSTS represents changeable aspects of the face, such as emotional expression, eye-gaze, and mouth movements, while ventral temporal fusiform cortex represents invariant aspects of faces, such as identity. This model was modified by O’Toole and colleagues (2002), who suggested that the rpSTS might also encode facial identity based on dynamic motion signatures. The pSTS face area (pSTS-FA) overlaps considerably with the pSTS region that represents biological motion (Engell & McCarthy, 2013; Grosbras, Beaton, & Eickhoff, 2012), and a causal role for the pSTS-FA in recognizing facial emotional expressions has been demonstrated using rTMS by Pitcher and colleagues (2014). In future work, it will be important to determine if this rpSTS region is comprised of a homogenous neuronal population, which codes emotional expression from a range of biological stimuli, or if there are separate neuronal populations tuned to different sources of information about the emotional state of another.

Dysfunction of the processing of biological motion in the pSTS region has been suggested to be a potential diagnostic ‘biomarker’ for social communication deficits in ASD (Björnsdotter et al., 2016), and it has been implicated in social deficits in schizophrenia (Kim et al., 2013). Our finding of a causal role of this region in emotion recognition from biological motion in healthy adults strengthens the rationale for examining rpSTS functionality when determining the basis of social and emotional deficits in clinical populations. In doing so, this work will provide a neurobiological basis to guide future treatment interventions for social and affective processing.

